# Interpretable Machine Learning and Comparative Genomics Reveal Microbial Plastic-Degrading (Microbeyt) Potential

**DOI:** 10.1101/2025.09.17.676701

**Authors:** Lokendra S. Thakur, Gurpreet Bharj, Manish Saroya

## Abstract

Plastic pollution poses a critical environmental threat, and microbial enzymes represent a sustainable strategy for polymer degradation. We present a computational pipeline that integrates orthogroup-based genomic analysis with machine learning and interpretable feature importance to identify microbial strains with high plastic-degrading potential. Using presence or absence matrices and SHAP-derived feature contributions to the MTP visualization, the workflow highlights conserved gene modules driving predictive classification. Application to a single genus revealed strains harboring versatile enzymatic repertoires capable of targeting diverse polymers, including polyethylene, polyethylene terephthalate, polyurethane, and polyhydroxyalkanoates. These findings provide a rational framework for prioritizing candidate strains for experimental validation and bioremediation strategies. Overall, this study demonstrates how integrating comparative genomics with interpretable machine learning can guide the systematic discovery of microbial solutions to plastic pollution.

## 1 Introduction

Plastic waste has become one of the most persistent environmental contaminants due to its durability and resistance to biodegradation. The world produces approximately 350 million tonnes of plastic waste annually, with an estimated 19 to 23 million tonnes leaking into aquatic ecosystems each year.**^1^** This alarming trend contributes to the accumulation of 75 to 199 million tonnes of plastic waste currently in our oceans.**^2^**

Despite efforts to mitigate this issue, only about 9.5% of the 400 million tonnes of plastic produced in 2022 was made from recycled materials, highlighting a significant reliance on fossil fuels for plastic production.**^3^** Furthermore, approximately 70% of plastic waste remains uncollected, leaks into the environment, is dumped into landfills, or subjected to open burning.**^4^**

While several bacterial species harbor enzymes capable of attacking synthetic polymers, the genomic determinants underlying these capabilities remain poorly characterized. Recent studies have begun to address this gap. Metagenomic analysis of plastic-degrading microbial communities has revealed novel microorganisms, metabolic pathways, and biocatalysts involved in polymer degradation.**^5^** Additionally, genomic mining has identified novel polyethylene terephthalate (PET)-degrading enzymes, although challenges remain in enhancing their efficiency for practical applications.**^6^**

The global plastics supply chain is highly complex, with trade-linked material flows amplifying environmental risks.**^7^** Microbial biodegradation has emerged as a promising avenue to mitigate these risks, converting plastic waste into valuable biochemical resources.**^8^**

This study aims to systematically analyze genomic conservation patterns within a single genus to identify candidate strains with high plastic-degrading potential. By leveraging advanced genomic tools and understanding the metabolic pathways involved, we can develop more efficient bioremediation strategies to combat the escalating plastic pollution crisis.

Our approach was inspired by the landmark study on the effector repertoire of *Legionella* species by Burstein et al.,**^9^** which used genome-wide ortholog clustering to reveal conserved functional modules. Building on this concept, we integrate orthogroup-based presence/absence matrices with machine learning and SHAP (SHapley Additive exPlanations)**^10^**-based feature importance to prioritize strains likely to possess versatile enzymatic repertoires. Publicly available genomic data (NCBI Entrez)**^11^** and curated knowledge of known plastic-degrading enzymes (*plasticDB* dataset^**12, 13**^) provide a reference framework for interpreting these predictions.

The resulting pipeline is modular, reproducible, and interpretable, offering a scalable approach to link conserved gene modules to predicted plastic-degrading potential. This framework can guide experimental validation and support the rational selection of microbial strains for bioremediation applications.

## 2 Data Selection and Genus Ranking

To identify the most promising candidates for plastic degradation, we analyzed the *plasticDB* dataset,^**12, 13**^ which catalogs microorganisms and enzymes involved in plastic biodegradation. This resource spans genus, species, and strain levels, providing a comprehensive view of microbial diversity linked to plastic breakdown. We prioritized species represented by many entries in the database, capturing diversity and experimental support, and exhibiting a broad repertoire of plastic-degrading enzymes, indicative of metabolic versatility. To ensure comparability across taxa, these species- and enzyme-level data were aggregated to the genus level, where scores were computed and genera were ranked accordingly.

Each genus was assigned a score derived from two weighted components. The first, the *species weight*, is proportional to the number of species reported for a genus. A larger species count suggests greater diversity and stronger experimental evidence, both of which increase the likelihood of identifying robust plastic degraders. The second, the *enzyme weight*, is proportional to the number of unique enzymes reported for the genus. A broader enzyme repertoire indicates the potential to degrade different polymer types or to cleave multiple chemical bonds within plastic substrates. As illustrated in Figure 1, the overall score was calculated as the sum of species and enzyme weights, with contributions normalized across the dataset to ensure comparability.

**Figure 1.**
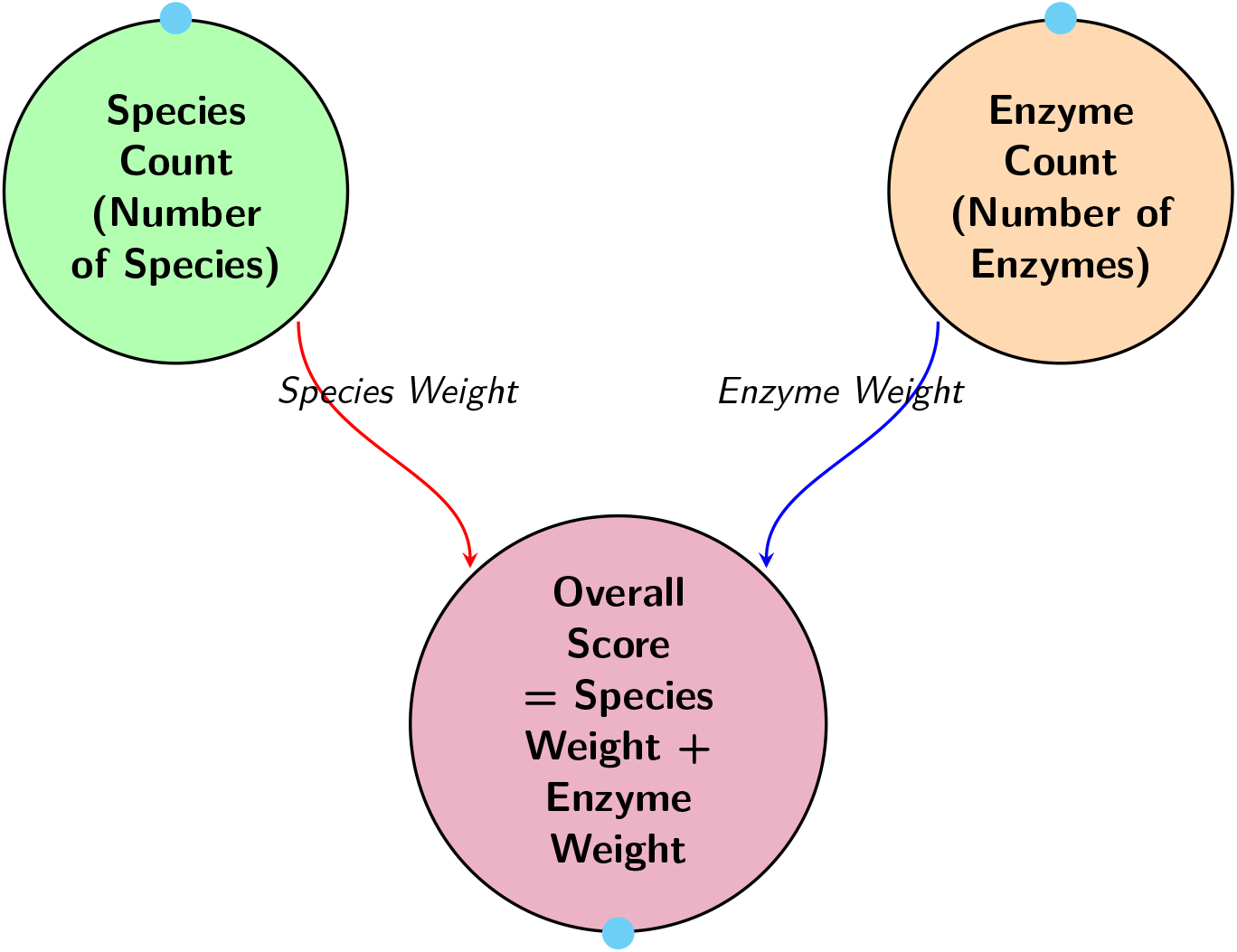
Visual representation of the scoring system. Species Count and Enzyme Count contribute jointly to the Overall Score.

Based on these computed scores, genera were ranked from highest to lowest. Table 1 lists the top 10 genera identified. Among them, *Pseudomonas* achieved the highest score due to its extensive species diversity and broad enzyme repertoire, followed by genera such as *Thermobifida, Cupriavidus*, and *Bacillus*, which showed strong potential but lacked the breadth observed in *Pseudomonas*.

**Table 1.** Top 10 genera from *plasticDB* ranked by score, based on species and enzyme representation.

| Genus | Rank | Species Count | Species Weight | Enzyme Count | Enzyme Weight | Score |
| --- | --- | --- | --- | --- | --- | --- |
| Pseudomonas | 1 | 25 | 0.1534 | 21 | 0.0399 | 0.1933 |
| Thermobifida | 2 | 13 | 0.0798 | 6 | 0.0114 | 0.0912 |
| Cupriavidus | 3 | 9 | 0.0552 | 3 | 0.0057 | 0.0609 |
| Bacillus | 4 | 5 | 0.0307 | 12 | 0.0228 | 0.0535 |
| Paucimonas | 5 | 6 | 0.0368 | 5 | 0.0095 | 0.0463 |
| Fusarium | 6 | 5 | 0.0307 | 7 | 0.0133 | 0.0440 |
| Rhodococcus | 7 | 4 | 0.0245 | 10 | 0.0190 | 0.0436 |
| Streptomyces | 8 | 4 | 0.0245 | 8 | 0.0152 | 0.0397 |
| Aspergillus | 9 | 2 | 0.0123 | 11 | 0.0209 | 0.0332 |
| Amycolatopsis | 10 | 3 | 0.0184 | 5 | 0.0095 | 0.0279 |

### 2.1 Data Curation and Input Dataset

After prioritizing genera, we focused on the top-ranked genus, *Pseudomonas*, for detailed strain-level analysis. For this purpose, we curated a dataset of 100 complete genomes from NCBI**^11^** RefSeq and GenBank (accession numbers listed in Supplementary Table S1). Only complete or high-quality assemblies were included to ensure reliable comparative genomics. Redundant entries were removed, and representative strains were selected to reflect the natural diversity of *Pseudomonas*, avoiding overrepresentation of any single species or environment.

The curated dataset includes both well-characterized species and unclassified strains labeled as *Pseudomonas sp*., providing a broad view of genomic diversity. Genome sizes ranged from 4.4 to 7.2 Mb, consistent with known variability within the genus and indicative of their metabolic adaptability. Notably, 31 strains were unclassified (*Pseudomonas sp*.), highlighting the extent of diversity yet to be fully resolved and representing potential sources of novel functional traits.

Figure 2 shows the distribution of the 100 curated *Pseudomonas* strains across 25 species and unclassified lineages. The chart demonstrates a clear imbalance in strain representation: a small number of species, such as *Pseudomonas sp*. (31 strains), *P. syringae* (26 strains), *P. aeruginosa* (9 strains), *P. fragariae* (7 strains), and *P. putida* (6 strains), collectively make up nearly 80% of the dataset. These taxa are either clinically significant or environmentally widespread, explaining the higher sequencing coverage.

**Figure 2.**
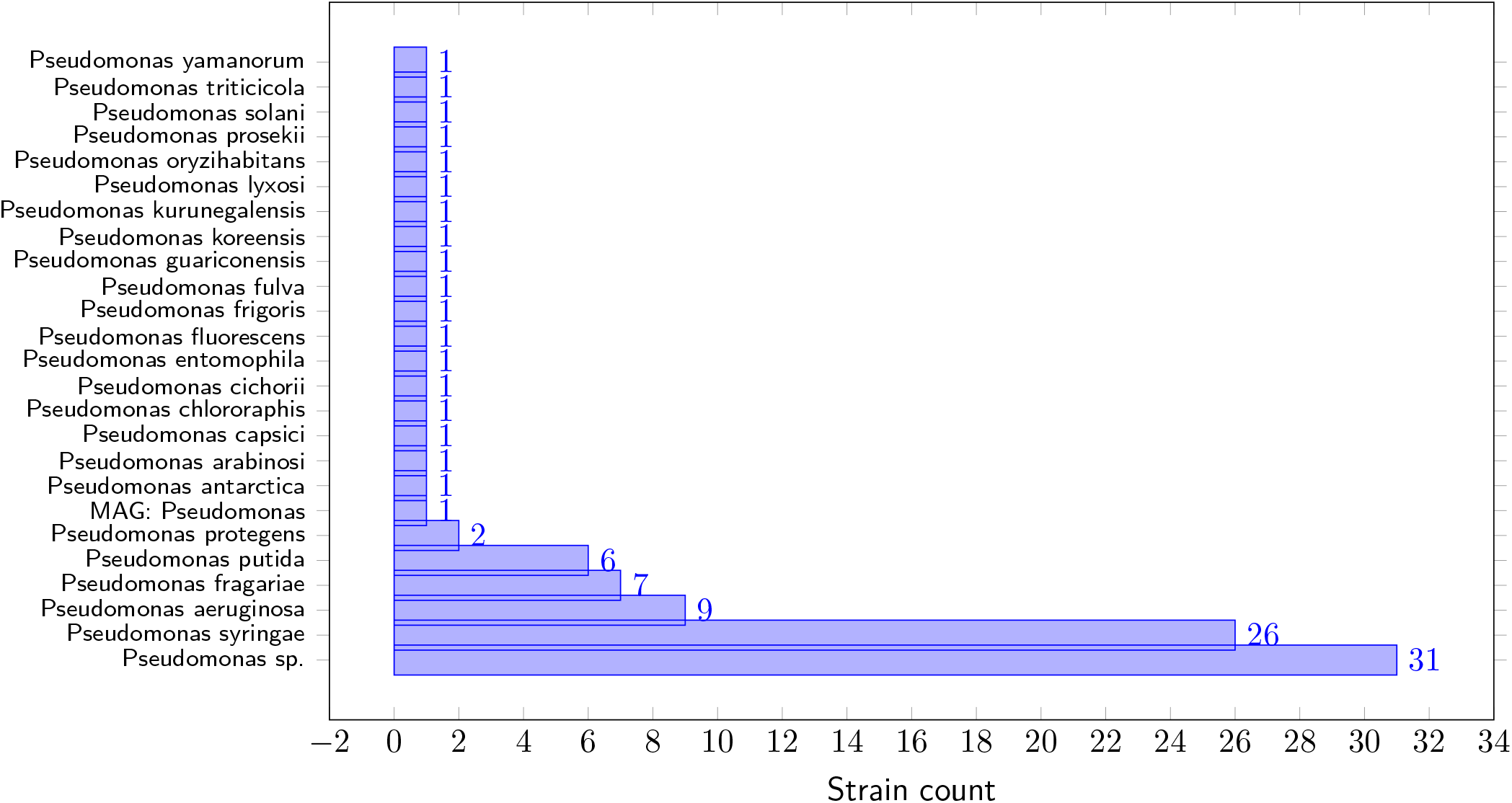
Distribution of the 100 curated *Pseudomonas* strains across 25 species. The chart highlights the predominance of a few species and the presence of many rare lineages represented by one or two strains each.

In contrast, the remaining 20 species are represented by only one or two strains each. This variability reflects both historical sequencing biases and the natural rarity of certain lineages. Including underrepresented species, despite low strain counts, is critical to capturing the full spectrum of *Pseudomonas* diversity. The presence of many single-strain species also suggests potential opportunities for discovering novel genes, metabolic pathways, or ecological adaptations that have not yet been extensively studied.

Overall, the bar chart highlights two key aspects of the curated dataset: (i) it preserves a robust representation of well-characterized species, allowing meaningful comparative genomics analyses, and (ii) it maintains taxonomic breadth by including rarer lineages, providing avenues for discovering previously uncharacterized genetic or functional diversity. By explicitly visualizing the number of strains per species, Figure 2 offers transparency into the dataset’s composition and informs the interpretation of downstream comparative analyses.

## 3 Methods

Building directly upon the data selection and genus ranking framework described in the previous section, where *Pseudomonas* was prioritized as the most promising genus due to its high representation of strains and enzyme diversity, we constructed a modular bioinformatics pipeline to predict and evaluate putative plastic-degrading effectors. The pipeline was implemented with a Makefile-based workflow to ensure reproducibility, automated dependency management, and seamless execution across multiple stages.

The overall design of the pipeline is illustrated in Figure 3, which is positioned alongside this section for direct reference.

**Figure 3.**
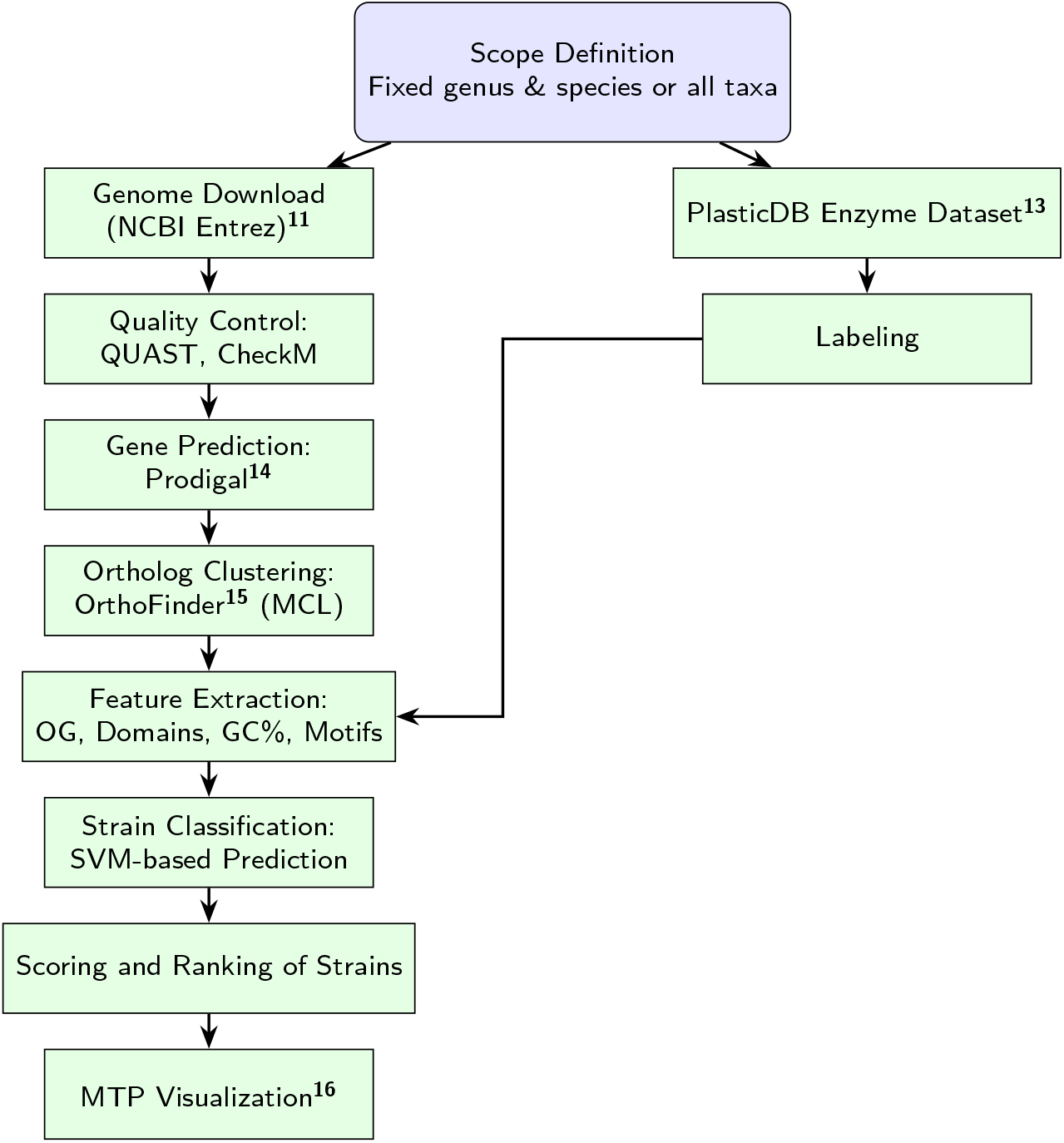
Pipeline for plastic degrading strain prediction integrating genome analysis with PlasticDB validation.

The pipeline begins with scope definition, where either a fixed genus and species or a broader set of taxa are selected based on the ranking system. For the current study, the genus *Pseudomonas* was chosen due to its dominance in both strain representation and enzyme diversity Table. 1. Genome assemblies were downloaded from NCBI Entrez,**^11^** ensuring comprehensive coverage across representative species. These assemblies were then subjected to quality assessment using QUAST (Quality Assessment Tool for Genome Assemblies)**^17^** to evaluate contiguity and assembly statistics, and CheckM**^18^** to estimate completeness and contamination, thereby ensuring reliable downstream predictions.

Following quality control, protein-coding genes were predicted using Prodigal,**^14^** a tool optimized for bacterial genomes, including incomplete drafts. The predicted proteomes were subsequently clustered into ortholog groups using OrthoFinder^**15, 19, 20**^ with Markov Clustering (MCL), which enabled the separation of conserved core proteins shared across all species from accessory proteins that may represent strain-specific innovations. This division was critical for distinguishing effectors with broad conservation from those confined to particular lineages.

### 3.1 Presence/Absence Matrix Construction

Orthogroups, as defined by OrthoFinder, represent sets of genes across bacterial genomes that trace back to a common ancestral gene, thereby capturing conserved evolutionary and functional relationships.

A gene belonging to an orthogroup indicates shared ancestry and potential similarity of function with its counterparts across other species. To translate this information into a clinically interpretable form, we constructed a **presence/absence matrix** (Figure. 4), where each row corresponds to a bacterial strain and each column corresponds to an orthogroup (OG). After preprocessing, this matrix had dimensions of **100 strains** × **41 orthogroups**, with each cell scored as “1” if the strain contained at least one gene from the orthogroup or “0” if it did not. To bridge genomic content with functional outcomes, we appended an additional column to this dataset containing binary phenotype labels from the PlasticDB resource, which indicate whether each strain has been experimentally associated with plastic degradation activity.

**Figure 4.**
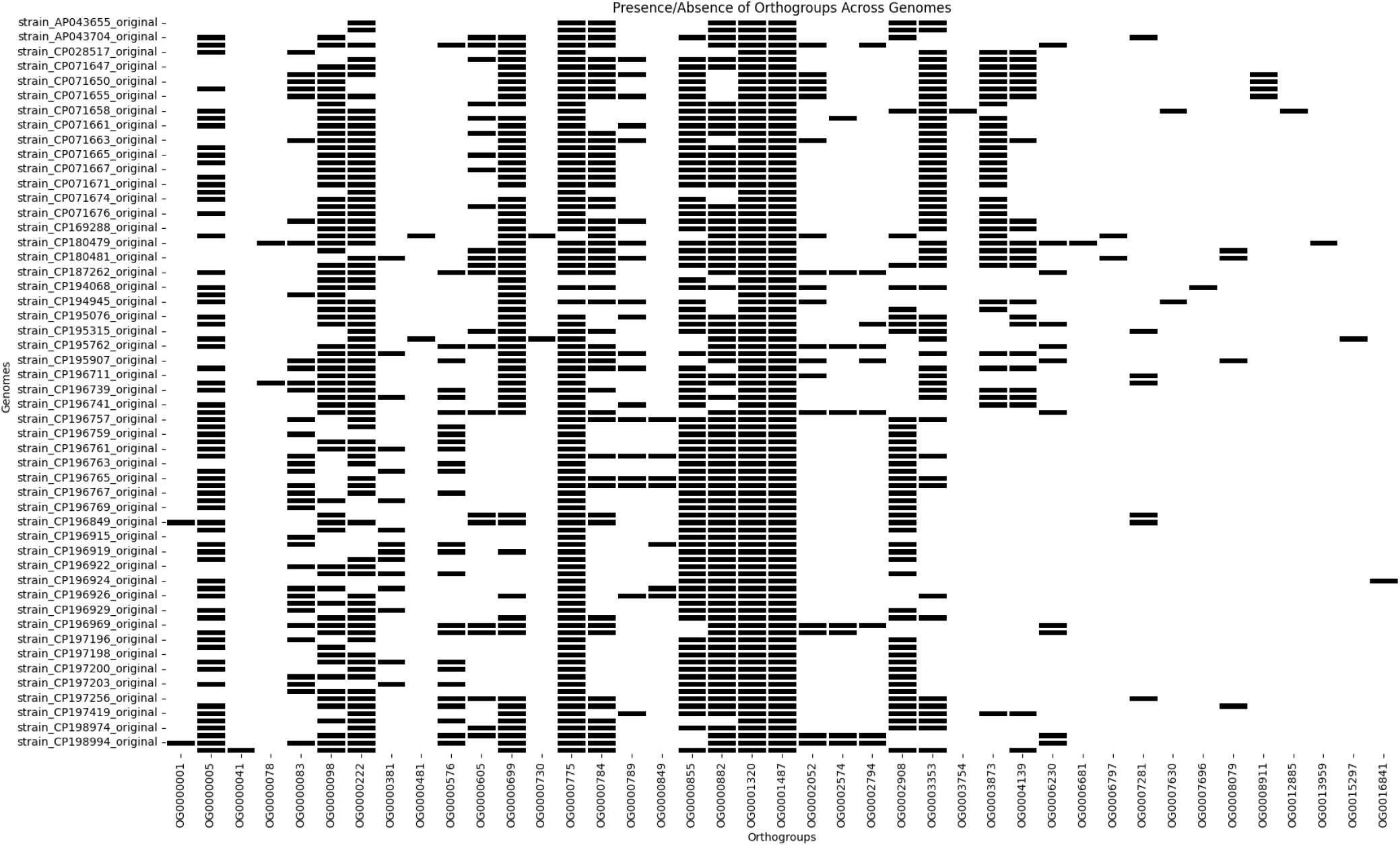
Presence/Absence of Orthogroups Across Genomes

The resulting integrated dataset was then used as the **input feature space for training a SVM (Support Vector Machine) classifier**, enabling systematic modeling of plastic-degrading potential from genomic content.

### 3.2 Visualization and Integration Framework

To interpret the contribution of individual orthogroups to classification outcomes, we employed a **Merkmal Treiber Plot (MTP)^16^** (Figure. 6). In this visualization, each colored wedge (arc) represents an orthogroup, with its angular width proportional to the number of strains (“carriers”) harboring that feature. Inner radial bands summarize the mean absolute SHAP (SHapley Additive exPlanations) values (Figure. 5) stratified by predicted degrader probability levels (Low, Medium, High), thereby linking orthogroup presence to model confidence. The outer black bars highlight the global importance of each orthogroup across all strains. Legends provide clinical and functional context by identifying the orthogroup, its representative species, the top strain carrying it. This visualization allowed us to prioritize orthogroups most strongly associated with biodegradation phenotypes, thereby offering both mechanistic interpretability of the classifier and actionable leads for identifying microbial strains with high translational potential in plastic waste remediation.

**Figure 5.**
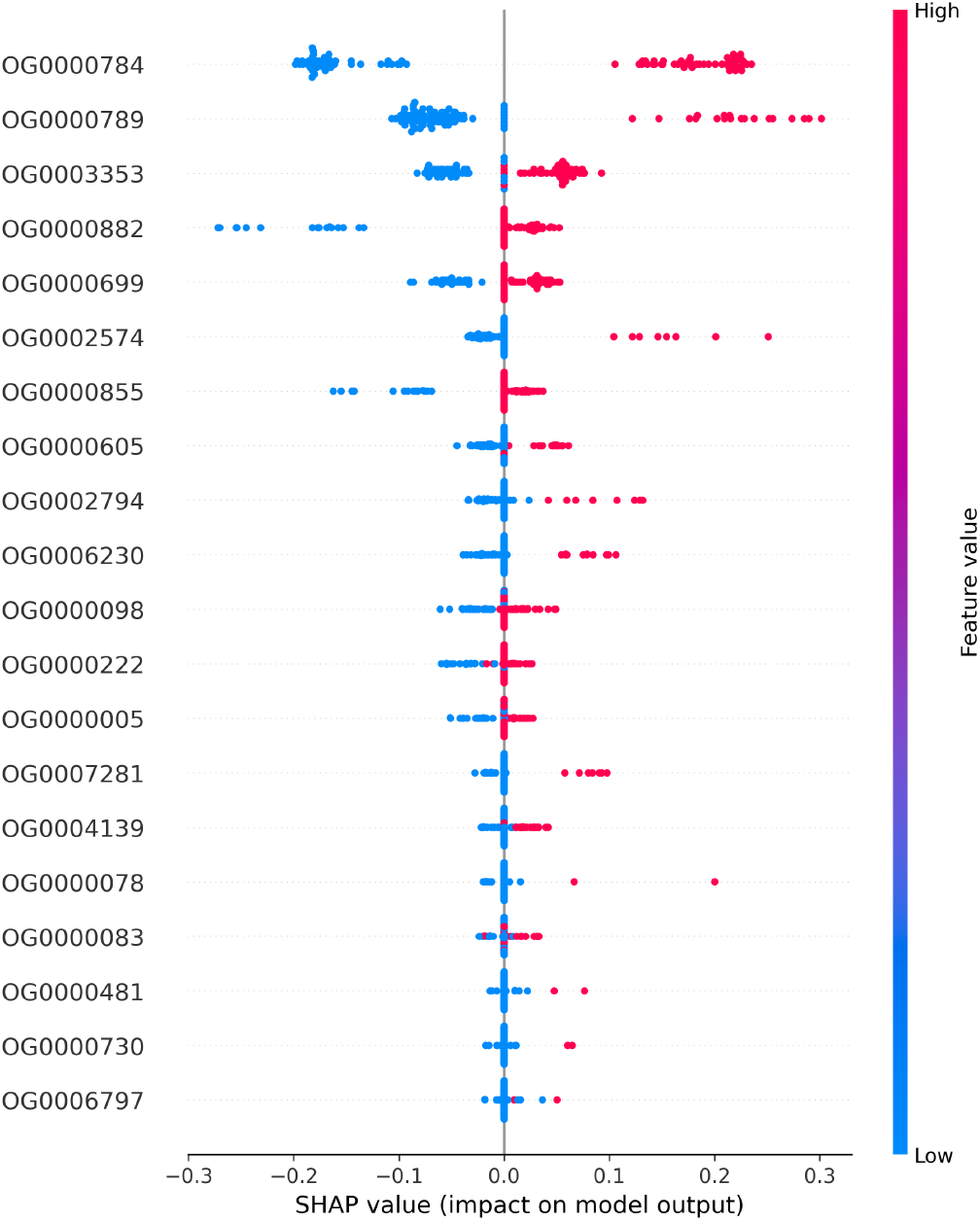
SHAP**^10^** top enriched Orthogroups

**Figure 6.**
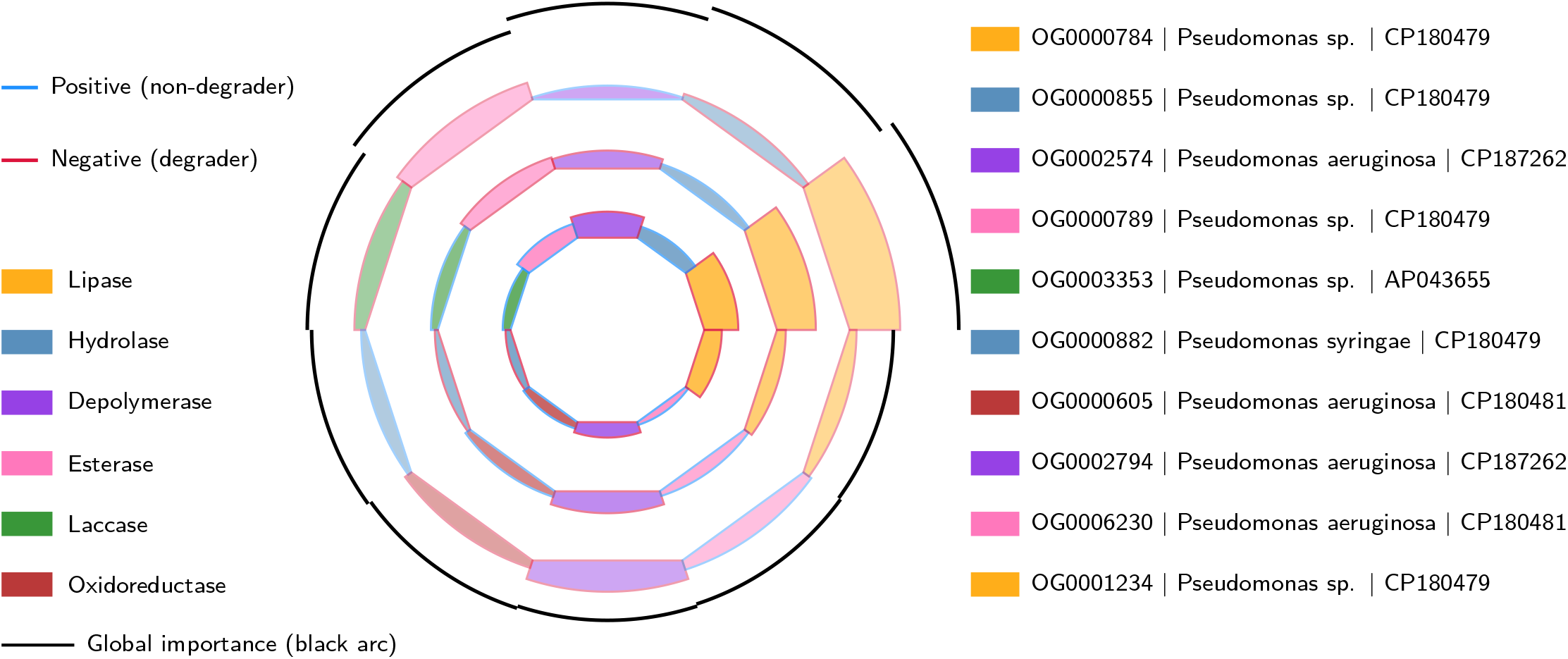
The MTP plot summarizes the contribution of orthogroups to plastic degradation across *Pseudomonas* strains. Each wedge corresponds to an orthogroup, with angular width showing how many strains carry it. Nested colored radial bands represent importance across “Low,” “Medium,” and “High” degrader groups: the fill color indicates enzyme functional class, while the outline color indicates whether the feature predicts degrader (red) or non-degrader (blue). The outer black arc shows global importance. This layered view allows rapid identification of which orthogroups, enzymes, and strains are most predictive for plastic degradation.

Finally, the distribution of **candidate degradative proteins** was examined in the context of the core and accessory genome framework. Core determinants represent conserved functions likely central to genus-wide survival, while accessory determinants highlight lineage-specific or niche-adaptive innovations. This combined strategy—integrating orthogroup-level modeling, protein-level annotation, and validation against PlasticDB—provides a comprehensive methodological framework for linking **candidate degrader strain** prioritization with protein-level discovery of degradation determinants, ultimately supporting the identification of microbial strains with high translational potential for plastic waste remediation.

## 4 Results

### Orthogroup Inference and Predictive Overview

Orthogroup analysis using OrthoFinder identified a diverse set of gene families across the Pseudomonas genomes included in our study. Among these, a subset of orthogroups was consistently enriched in strains associated with plastic degradation potential.

Predictive modeling, evaluated through true and predicted class assignments with associated probabilities, showed high accuracy in distinguishing degrading from non-degrading strains. For example, Pseudomonas strain CP180481 was predicted with a probability of 0.99 to be plastic-degrading, strongly supporting its enzymatic repertoire. Overall, the model demonstrated robust performance, with most positive strains correctly classified and only a limited number of false negatives (e.g., strain CP071658 with a predicted probability of 0.23 despite its known degrading potential).

### Core and Accessory Orthogroups Relevant to Plastic Degradation

From the inferred orthogroups, those associated with enzymes such as hydrolases, esterases, and depolymerases were disproportionately represented in predicted degraders. OG0000784 emerged as the most important predictor with the highest mean absolute SHAP**^10^** value, suggesting that its encoded functions play a central role in plastic biodegradation. The presence of core orthogroups across multiple Pseudomonas species indicates shared metabolic strategies, while accessory orthogroups contribute to strain-specific versatility.

This mirrors evolutionary trends observed in other metabolic systems, where core functions provide baseline activity and accessory functions extend substrate specificity.

### Functional Annotation of Enzyme Families

Annotation of the enriched orthogroups revealed a broad enzymatic spectrum linked to different classes of plastics. Polyhydroxyalkanoate (PHA) and polyhydroxybutyrate (PHB) depolymerases were abundant, reflecting natural adaptation of Pseudomonas to aliphatic polyesters. Cutinases and esterases were linked to polyethylene terephthalate (PET) and polybutylene succinate adipate (PBSA) degradation, while laccases and hydrolases were associated with polyethylene (PE), polystyrene (PS), and polyvinyl chloride (PVC). Polyurethane (PU)-active enzymes, including polyurethane esterases and Impranil-degrading esterases, were detected in specific lineages. This diversity indicates that plastic-degrading Pseudomonas strains rely on a distributed enzymatic toolkit rather than a single universal pathway.

### Strain-Level Contributions and Key Predictors

At the strain level, Pseudomonas sp. CP180479 stood out as the top contributor across multiple orthogroups, reflecting its broad degradation potential. Other strongly predicted degraders included strains **CP180480, CP180481, and CP187262**, each supported by high predicted probabilities (*>* 0.87). Conversely, strains such as **CP071658** and **CP071652** were misclassified as non-degraders, likely reflecting either incomplete genome annotations or underrepresentation of accessory orthogroups in the training dataset. Because accessory orthogroups often encode strain-specific functions, missing or partial annotations can obscure their contribution to degradation potential. In contrast, core orthogroups—being widely conserved—are more robust to minor annotation gaps. Nonetheless, predictive accuracy generally requires near-complete genome assemblies, with annotation completeness of ≥95% considered optimal,^**21, 22**^ and values below 90% carrying substantial risk of misclassification.^**21, 22**^ These results emphasize the importance of both core orthogroups and strain-specific variations in shaping the predictive landscape, while also emphasizing the need for high-quality, complete genome annotations in such analyses.

### Model Feature Importance and SHAP Analysis

To better understand the contributions of individual features, we employed SHAP (SHapley Additive exPlanations)**^10^** analysis. The highest-scoring orthogroups, led by OG0000784, 0G0000789, and 0G0003353, were enriched for hydrolases and depolymerases, enzymes directly implicated in polymer breakdown. The distribution of mean absolute SHAP values confirmed that only a small subset of orthogroups explained the majority of predictive performance, highlighting potential marker genes for future biodegradation screening. This hierarchical importance structure aligns with ecological expectations, where a few key enzymes mediate most of the degradation activity, supported by peripheral accessory functions.

### Integrative Flow of Results

Taken together, these findings indicate that Pseudomonas genomes encode a mixture of core and accessory orthogroups that collectively drive plastic degradation. Core sets provide broad-spectrum depolymerase activity, while accessory sets fine-tune substrate range toward PET, PU, PS, PVC, Nylon, and other polymers. The predictive framework, validated by strong agreement between true and predicted degraders, highlights a limited but powerful set of orthogroups as the central drivers of plastic biodegradation potential in Pseudomonas. These results complement the data and methodological framework described earlier, demonstrating that orthogroup-centered approaches can both classify degraders with high precision and uncover the enzymatic basis of plastic degradation diversity.

### 4.1 Merkmal Treiber Plot (MTP) Analysis: Top Microbeyts (Plastic-Degrading Microbes)

To complement the pipeline described earlier, we built the MTP**^16^** framework in order to visualize and interpret which orthogroups most strongly predict plastic degradation across *Pseudomonas* strains. Features (orthogroups) were ranked by mean absolute SHAP values, then paired with strain-metadata (species, enzyme annotations, plastics degraded) to generate the MTP, which reveals both the breadth and strength of each feature. In the plot, the **circle is divided into wedges (sectors)**, each corresponding to an orthogroup. The **angular width** of a wedge shows how many strains carry that orthogroup. **Nested radial colored bands** within each wedge display the contribution of that orthogroup across “Low,” “Medium,” and “High” degrader probability groups: the **fill color** indicates the enzyme functional class (e.g., lipase, hydrolase, esterase), while the **band outline color** indicates whether the feature pushes the prediction toward degrader (positive, red) or non-degrader (negative, blue). Finally, the **outer black arc** along the edge of each wedge depicts the global importance of that orthogroup across all strains. Each wedge is labeled “OG | Species | Top-strain,” where OG is the orthogroup, Species is the species carrying that orthogroup, and Top-strain is the carrier strain with the highest model probability.

From this MTP analysis, several strongly predictive orthogroups emerged, with actual enzyme–plastic pairings supporting their mechanistic relevance. For example, the orthogroup OG0000784 is associated with *Pseudomonas sp*., top strain CP180479, and is linked with enzymes such as lipase, PHA depolymerase, PHB depolymerase and oxidized PVA hydrolase, which in turn correspond to plastics including HDPE,**^23^** PET, PU,**^24^** PBSA,**^25^** PE,^**26**–**29**^ LDPE**^30^** and PS.**^31^** The broad substrate range reflected by those plastics mirrors the high importance of that OG in both global and high-probability degrader clusters. Another orthogroup, OG0000855, also associated with *Pseudomonas sp*., shows enzymes like polyurethanase, esterases, and hydrolase, and plastics such as PU, LDPE, and HDPE. While its global importance is lower, the MTP plot shows that in “High degrader” probability strains, it contributes conspicuously, suggesting a strong role in strains highly capable of plastic degradation. A further example, OG0002574 in *Pseudomonas aeruginosa*, carries esterase, lipase, and PHB depolymerase annotations, and correlates with plastics including PET,**^32^** PU,**^33^** PE,^**29, 34**^ and PE blends. Although this OG has moderate global SHAP value, its presence in strain CP187262 and alignment with these enzyme–plastic pairs emphasize its value as a functional marker.

The MTP plot thus offers layered insight: it does not merely rank features by statistical importance but integrates biological annotation (which enzymes are involved, which plastics are targeted), directional predictive signal (toward degrader vs. non-degrader), and strain-level metadata (which strain carries that OG most prominently). For clinicians or biotechnologists interested in selecting candidate strains or enzyme systems for experimental validation or bioremediation, MTP provides a guide to which orthogroups may be most promising both globally and in high-activity contexts.

## 5 Discussion

Our study identified top-predicted microbial strains with high plastic-degrading potential through the integration of orthogroup (OG) clustering and SHAP-based feature importance analysis. Orthogroups represent sets of homologous genes shared across strains, and their presence in specific strains indicates conserved genetic modules potentially linked to function. By applying SHAP analysis, we quantified the contribution of each OG to the model’s prediction, thereby highlighting which gene clusters most strongly drive the classification of a strain as a plastic degrader. Strains enriched in high-impact OGs therefore represent prime candidates for further experimental validation, as their genomic content is most predictive of broad enzymatic capabilities against synthetic polymers. Among the top-predicted lineages, *Pseudomonas sp*. emerged consistently across multiple OGs. In OG0000784, *CP180479* was identified as the top strain, representing an OG encompassing 72 strains collected from soil, freshwater, wastewater, and industrial sites. SHAP-ranked OGs in this strain encode enzymes including alkane hydroxylase, esterase, hydrolase, laccase, lipase, PHB depolymerase, PHA depolymerase, polyurethanase, oxidized PVA hydrolase, and PVA dehydrogenase, reflecting a remarkably versatile enzymatic repertoire. These enzymes target a wide variety of plastics, such as HDPE, LDPE, PU, PCL, PET, PHB, PHA, PLA, PS, PVC, O-PVA, and Nylon, highlighting the broad polymer-degrading potential of this lineage. Similarly, OG0000789 and OG0003353 also featured *Pseudomonas sp. CP180479* and *AP043655* as top representatives, with 63 and 52 strains in their respective OGs. OG0000882, led by *Pseudomonas syringae CP180479*, included 68 member strains collected from soil, rhizosphere, and aquatic ecosystems. These top-predicted OGs encode overlapping sets of enzymes, enabling degradation of HDPE, LDPE, PET, PHA, PHB, PU, PVA, PVC, and related polymer blends. The member strains were isolated from diverse environments including soil, marine sediments, landfills, sewage sludge, and polluted intertidal regions, spanning countries such as Iran, India, USA, Taiwan, Pakistan, China, Japan, South Korea, Nigeria, Thailand, Svalbard, and Serbia. *Pseudomonas aeruginosa* appeared prominently in OG0002574, OG0000605, OG0002794, and OG0006230, with *CP180481* and *CP187262* serving as the top representatives. These OGs comprised 9, 23, 10, and 11 member strains respectively, originating from landfills, soils, dumping sites, and culture collections across India, Japan, South Korea, and Iran. The top OGs in these strains encode alkane hydroxylase, esterase, hydrolase, laccase, lipase, PHB depolymerase, PVA dehydrogenase, and various polyurethanases, facilitating degradation of plastics including HDPE, LDPE, PU, PBSA, PHB, PET, PS, and PU blends.

The observed distribution of top-predicted species and strains illustrates the polygenic nature of plastic degradation, with multiple enzyme classes acting synergistically across diverse polymer backbones. Membership in each OG implies that all strains share conserved genetic modules, suggesting potential for plastic-degrading function even in less-characterized strains. The combination of orthogroup conservation and SHAP-driven feature importance provides an interpretable framework linking genomic content to predicted plastic-degrading function. The ecological and geographic diversity of these strains, coupled with the number of strains per OG, emphasizes the importance of environmental context in shaping degradative capabilities and supports their selection for laboratory validation, bioremediation trials, and the rational design of microbial consortia for targeted polymer breakdown in polluted environments.

Details of all strains within each top-predicted OG, are provided in Supplementary Table S2.

## 6 Limitations

This study has limitations that should be considered when interpreting the results. First, presence/absence matrices rely on accurate orthogroup assignments, which can be influenced by genome completeness and annotation quality within the genus. Second, limited representation of certain clades may reduce clustering resolution and affect the robustness of inferred relationships. Third, the analysis captures genomic potential rather than experimentally confirmed functions, so observed patterns may not directly translate to phenotypic traits. Finally, as this study is focused on a single genus, the findings and conclusions may not be generalizable to other bacterial taxa.

## 7 Conclusion and Future Work

This study presents a modular and interpretable pipeline for systematically linking genomic content to predicted plastic-degrading potential in microbial strains. By integrating orthogroup clustering with SHAP-based feature importance analysis, the framework identifies conserved gene modules that contribute most strongly to predictive models, highlighting strains with the highest likelihood of enzymatic activity against diverse plastics. The combination of presence/absence matrices, clustering, and interpretable model outputs provides a transparent and reproducible approach for prioritizing candidate strains for experimental validation.

Application of this pipeline to a single genus revealed biologically meaningful patterns, with top-predicted strains such as *Pseudomonas sp*. consistently harboring orthogroups encoding enzymes capable of degrading a broad spectrum of polymers, including HDPE, LDPE, PET, PU, PHA, PHB, PLA, PS, PVC, and PVA. The analysis demonstrated the polygenic nature of plastic degradation, where multiple enzyme classes act synergistically, and conserved orthogroups allow inference of degradative potential even in less-characterized strains. The geographic and ecological diversity of these strains further emphasizes the importance of environmental context in shaping enzymatic capabilities.

### Future Work

Several directions can extend the biological and computational scope of this work. Experimental validation of top-ranked strains and orthogroups remains a priority to confirm predicted enzymatic activity. Expanding the pipeline to additional strains within the genus and closely related taxa will improve the resolution of clustering and enhance generalizability. Future annotation enrichment should leverage tools such as **InterProScan**^**35**–**37**^ and **Pfam^38^** to systematically identify conserved protein domains, motifs, and catalytic sites, enabling the derivation of multidimensional feature sets—including amino acid composition, GC content, and domain architectures—for more precise functional inference. Incorporating structural modeling and pathway-level annotation (e.g., KEGG, PlasticDB) could further refine predictions and provide mechanistic insight into substrate specificity. Ensemble learning approaches may improve predictive accuracy, reducing uncertainty in orthogroup-based inference. Finally, development of interactive visualizations will facilitate exploration of strain-specific predictions, orthogroup conservation patterns, and SHAP-derived feature importance, supporting translational applications in environmental biotechnology and microbial ecology.

### Final Remarks

By systematically combining orthogroup clustering, interpretable predictive modeling, and curated annotation, the pipeline offers a robust framework for discovering microbial strains and gene modules with potential plastic-degrading activity. This approach bridges computational prediction with biological insight, enabling high-throughput prioritization of candidate strains and providing actionable guidance for experimental validation, bioremediation strategies, and the rational design of microbial consortia for polymer degradation.

## Acknowledgements

Data used in the preparation of this article were obtained from publicly available nucleotide and protein sequences in the **National Center for Biotechnology Information (NCBI) RefSeq** and **GenBank^39^** databases. RefSeq provides curated, non-redundant reference sequences,**^40^** and GenBank contains publicly submitted sequences.**^41^** Accession numbers for all sequences used are provided in Supplementary Table S1.

We are grateful to Dr. Eyal Y. Kimchi (Department of Neurology, Northwestern University Feinberg School of Medicine, USA), Dr. Strajit Ghosh (Massachusetts General Hospital and Harvard Medical School, USA), Dr. Fernando Gómez-Baquero (Jacobs Technion–Cornell Institute, Cornell Tech; NSF Upstate New York Energy Storage Engine, USA), Dr. Kamana Porwal (Department of Mathematics, IIT Delhi, India), and Dr. Mustafa Hajij (MSDSAI Program, University of San Francisco, California, USA) for their valuable guidance, support, and insightful suggestions, which greatly contributed to this work.

## Author Contributions

-Dr. Lokendra S. Thakur conceived the idea, designed the study, developed method and algorithm, curated data and done analysis, developed pipeline, all sections writing. -Dr. Gurpreet Bharj performed the microbiological and clinical analysis. -Manish Saroya checked citations and contributed in refining content. -All authors reviewed and approved the final version.

## Ethics Approval and Consent to Participate

Not applicable.

## Data Availability

We accessed raw data from the publicly available repository: https://www.ncbi.nlm.nih.gov/ Code supporting this study are available on reasonable request.

## Disclaimer

Preprints are preliminary reports that have not been peer reviewed. They should not be regarded as conclusive, guide clinical practice, or be reported in news media as established information.

## Supplementary

**Supplementary Table S1.** Curated *Pseudomonas* genome dataset from NCBI RefSeq/GenBank.

| Accession | Species | Status | Message |
| --- | --- | --- | --- |
| CP198974 | *Pseudomonas kurunegalensis* | Success | 5855340 bp |
| CP198993 | *Pseudomonas aeruginosa* | Success | 6522294 bp |
| AP043656 | *Pseudomonas cichorii* | Success | 5918927 bp |
| CP197419 | *Pseudomonas chlororaphis* | Success | 7052953 bp |
| AP043655 | *Pseudomonas capsici* | Success | 4852321 bp |
| CP196970 | *Pseudomonas aeruginosa* | Success | 6551601 bp |
| CP169288 | *Pseudomonas* sp. | Success | 7205777 bp |
| CP199001 | *Pseudomonas* sp. | Success | 6006889 bp |
| CP197196 | *Pseudomonas fragariae* | Success | 5887932 bp |
| CP198994 | *Pseudomonas aeruginosa* | Success | 6344686 bp |
| CP195082 | *Pseudomonas protegens* | Success | 7233958 bp |
| CP197201 | *Pseudomonas syringae* | Success | 5861675 bp |
| CP197353 | *Pseudomonas* sp. | Success | 6069357 bp |
| AP043704 | *Pseudomonas putida* | Success | 6183974 bp |
| CP197256 | *Pseudomonas putida* | Success | 6032578 bp |
| CP197197 | *Pseudomonas syringae* | Success | 5839168 bp |
| CP198970 | *Pseudomonas* sp. | Success | 7255948 bp |
| CP028517 | *Pseudomonas fluorescens* | Success | 6515198 bp |
| CP197204 | *Pseudomonas syringae* | Success | 5917844 bp |
| CP197203 | *Pseudomonas syringae* | Success | 5879092 bp |
| CP197199 | *Pseudomonas syringae* | Success | 5484274 bp |
| CP197198 | *Pseudomonas syringae* | Success | 5992041 bp |
| CP197200 | *Pseudomonas syringae* | Success | 5448011 bp |
| CP178184 | *Pseudomonas* sp. | Success | 6719236 bp |
| CP196925 | *Pseudomonas fragariae* | Success | 5955777 bp |
| CP196927 | *Pseudomonas syringae* | Success | 5893211 bp |
| CP196922 | *Pseudomonas syringae* | Success | 5927426 bp |
| CP196969 | *Pseudomonas aeruginosa* | Success | 6578177 bp |
| CP196913 | *Pseudomonas syringae* | Success | 5950303 bp |
| CP196923 | *Pseudomonas fragariae* | Success | 5956165 bp |
| CP196915 | *Pseudomonas fragariae* | Success | 6146788 bp |
| CP196924 | *Pseudomonas fragariae* | Success | 6141266 bp |
| CP196929 | *Pseudomonas syringae* | Success | 5529192 bp |
| CP196920 | *Pseudomonas syringae* | Success | 5969903 bp |
| CP196749 | *Pseudomonas aeruginosa* | Success | 6589538 bp |
| CP196768 | *Pseudomonas syringae* | Success | 5577486 bp |
| CP196919 | *Pseudomonas syringae* | Success | 5980508 bp |
| CP196917 | *Pseudomonas fragariae* | Success | 6065732 bp |
| CP196767 | *Pseudomonas syringae* | Success | 6014304 bp |
| CP196761 | *Pseudomonas syringae* | Success | 5607559 bp |
| CP196763 | *Pseudomonas syringae* | Success | 5992788 bp |
| CP196764 | *Pseudomonas syringae* | Success | 5807824 bp |
| CP196766 | *Pseudomonas syringae* | Success | 6014147 bp |
| CP196759 | *Pseudomonas syringae* | Success | 6024633 bp |
| CP196926 | *Pseudomonas fragariae* | Success | 6012559 bp |
| CP196958 | *Pseudomonas putida* | Success | 5768295 bp |
| CP196769 | *Pseudomonas syringae* | Success | 5581818 bp |
| CP196758 | *Pseudomonas syringae* | Success | 5807276 bp |
| CP196849 | *Pseudomonas putida* | Success | 5907346 bp |
| CP196848 | *Pseudomonas putida* | Success | 5909934 bp |
| CP196765 | *Pseudomonas syringae* | Success | 5989840 bp |
| CP196711 | *Pseudomonas* sp. | Success | 4498643 bp |
| CP196762 | *Pseudomonas syringae* | Success | 6241719 bp |
| CP196713 | *Pseudomonas* sp. | Success | 4815009 bp |
| CP196760 | *Pseudomonas syringae* | Success | 5546732 bp |
| CP187262 | *Pseudomonas aeruginosa* | Success | 6326572 bp |
| CP196757 | *Pseudomonas syringae* | Success | 6123328 bp |
| CP196740 | *Pseudomonas yamanorum* | Success | 7078232 bp |
| CP194068 | *Pseudomonas guariconensis* | Success | 5759654 bp |
| CP196739 | *Pseudomonas prosekii* | Success | 6090943 bp |
| CP196741 | *Pseudomonas antarctica* | Success | 6084575 bp |
| CP195907 | *Pseudomonas aeruginosa* | Success | 6961963 bp |
| CP196684 | *Pseudomonas koreensis* | Success | 6181985 bp |
| CP180479 | *Pseudomonas frigoris* | Success | 6805123 bp |
| CP195831 | *Pseudomonas* sp. | Success | 6330929 bp |
| CP191382 | *Pseudomonas fulva* | Success | 4897385 bp |
| CP195762 | *Pseudomonas aeruginosa* | Success | 6592422 bp |
| CP180481 | *Pseudomonas lyxosi* | Success | 6722390 bp |
| CP020603 | *Pseudomonas aeruginosa* | Success | 7040952 bp |
| CP180480 | *Pseudomonas arabinosi* | Success | 6932784 bp |
| CP136718 | *Pseudomonas* sp. | Success | 6452033 bp |
| CP182239 | MAG: *Pseudomonas* | Success | 6101023 bp |
| CP071676 | *Pseudomonas* sp. | Success | 5929390 bp |
| CP194941 | *Pseudomonas oryzihabitans* | Success | 4785578 bp |
| CP194945 | *Pseudomonas triticicola* | Success | 6005210 bp |
| CP071675 | *Pseudomonas* sp. | Success | 5617489 bp |
| CP071674 | *Pseudomonas* sp. | Success | 5667114 bp |
| CP194950 | *Pseudomonas entomophila* | Success | 4461257 bp |
| CP071673 | *Pseudomonas* sp. | Success | 4980928 bp |
| CP071667 | *Pseudomonas* sp. | Success | 5617520 bp |
| CP195315 | *Pseudomonas putida* | Success | 5844059 bp |
| CP071666 | *Pseudomonas* sp. | Success | 5625150 bp |
| CP195316 | *Pseudomonas solani* | Success | 6676092 bp |
| CP195076 | *Pseudomonas protegens* | Success | 7053038 bp |
| CP071668 | *Pseudomonas* sp. | Success | 5764948 bp |
| CP071665 | *Pseudomonas* sp. | Success | 5740047 bp |
| CP071671 | *Pseudomonas* sp. | Success | 5625157 bp |
| CP071663 | *Pseudomonas* sp. | Success | 6151268 bp |
| CP071652 | *Pseudomonas* sp. | Success | 6118271 bp |
| CP071662 | *Pseudomonas* sp. | Success | 5566958 bp |
| CP071658 | *Pseudomonas* sp. | Success | 4889755 bp |
| CP071664 | *Pseudomonas* sp. | Success | 5625203 bp |
| CP071649 | *Pseudomonas* sp. | Success | 6118252 bp |
| CP071644 | *Pseudomonas* sp. | Success | 6931860 bp |
| CP071647 | *Pseudomonas* sp. | Success | 6850403 bp |
| CP071661 | *Pseudomonas* sp. | Success | 5982333 bp |
| CP071660 | *Pseudomonas* sp. | Success | 6078262 bp |
| CP071657 | *Pseudomonas* sp. | Success | 6150913 bp |
| CP071650 | *Pseudomonas* sp. | Success | 6118259 bp |
| CP071655 | *Pseudomonas* sp. | Success | 6118238 bp |

**Supplementary Table S2.** Features, strains, enzymes, plastics, and SHAP values identified in *Pseudomonas species*.

| Feature | Species | Top Strain | All Strains | Enzymes | Plastics | Mean Abs SHAP |
| --- | --- | --- | --- | --- | --- | --- |
| OG0000784 | *Pseudomonas* sp. | CP180479 | AP043655, AP043656, AP043704, CP020603, CP071644, CP071647, CP071649, CP071650, CP071652, CP071655, CP071662, CP071663, CP071664, CP071665, CP071666, CP071667, CP071668, CP071671, CP071674, CP071675, CP136718, CP169288, CP178184, **CP180479**, CP180480, CP180481, CP182239, CP187262, CP194068, CP194945, CP195315, CP195762, CP195831, CP195907, CP196684, CP196739, CP196749, CP196757, CP196762, CP196765, CP196766, CP196848, CP196849, CP196958, CP196969, CP196970, CP197256, CP197353, CP198970, CP198974, CP198993, CP198994, CP199001 | Alkane hydroxylase, Esterase, Hydrolase, Laccase, Lipase, No, Oxidized PVA hydrolase, PHA depolymerase, PHB depolymerase, PVA dehydrogenase, Polyurethanase | HDPE, Impranil, LDPE, Nylon, O-PVA, PBS, PBSA, PCL, PE, PEG, PES, PET, PHA, PHA Blend, PHB, PHN, PHO, PLA, PP, PS, PS Blend, PU, PU Blend, PVA, PVC, PVC Blend | 0.17644 |
| OG0000789 | *Pseudomonas* sp. | CP180479 | CP071644, CP071649, CP071655, CP071661, CP071663, CP136718, **CP180479**, CP180481, CP194945, CP195076, CP195831, CP196684, CP196741, CP196757, CP196762, CP196765, CP196766, CP196926, CP197419 | Alkane hydroxylase, Esterase, Hydrolase, Lipase, No, Oxidized PVA hydrolase, PHA depolymerase, PVA dehydrogenase, Polyurethanase, Polyurethanase A, Polyurethanase B, Polyurethanase esterase, Polyurethane esterase A, Serine hydrolase | HDPE, Impranil, LDPE, Nylon, O-PVA, PBS, PBSA, PCL, PE, PEG, PES, PET, PHA, PHA Blend, PHB, PHBV, PHN, PHO, PLA, PP, PS, PS Blend, PU, PVA, PVC | 0.09390 |
| OG0003353 | *Pseudomonas* sp. | CP180479 | AP043655, AP043656, CP028517, CP071644, CP071647, CP071649, CP071650, CP071652, CP071655, CP071657, CP071658, CP071660, CP071661, CP071662, CP071663, CP071664, CP071665, CP071666, CP071667, CP071668, CP071671, CP071673, CP071674, CP071675, CP071676, CP136718, CP169288, **CP180479**, CP180480, CP180481, CP182239, CP194068, CP195076, CP195082, CP195315, CP195316, CP195831, CP196684, CP196711, CP196713, CP196739, CP196740, CP196757, CP196762, CP196765, CP196766, CP196926, CP196958, CP197256, CP197353, CP197419, CP198970, CP198974, CP199001 | Alkane hydroxylase, Esterase, Hydrolase, Lipase, No, Oxidized PVA hydrolase, PHA depolymerase, PHB depolymerase, PVA dehydrogenase, Polyurethanase, Polyurethanase A, Polyurethanase B, Polyurethanase esterase, Polyurethane esterase A, Protease, Serine hydrolase | HDPE, Impranil, LDPE, Nylon, O-PVA, PBS, PBSA, PCL, PE, PEG, PES, PET, PHA, PHA Blend, PHB, PHBV, PHN, PHO, PLA, PP, PS, PS Blend, PU, PVA, PVC | 0.04973 |
| OG0000882 | *Pseudomonas syringae* | CP180479 | AP043655, AP043656, AP043704, CP020603, CP071644, CP071647, CP071657, CP071658, CP071660, CP071661, CP071662, CP071664, CP071665, CP071666, CP071667, CP071668, CP071671, CP071675, CP071676, CP136718, CP169288, CP178184, **CP180479**, CP180480, CP180481, CP182239, CP187262, CP194068, CP195076, CP195082, CP195315, CP195316, CP195762, CP195831, CP195907, CP196711, CP196713, CP196740, CP196749, CP196757, CP196758, CP196759, CP196760, CP196761, CP196762, CP196763, CP196764, CP196765, CP196766, CP196767, CP196768, CP196769, CP196848, CP196849, CP196913, CP196915, CP196917, CP196919, CP196920, CP196922, CP196923, CP196924, CP196925, CP196926, CP196927, CP196929, CP196958, CP196969, CP196970, CP197196, CP197197, CP197198, CP197199, CP197200, CP197201, CP197203, CP197204, CP197256, CP197353, CP197419, CP198974, CP198993, CP198994, CP199001 | Alkane hydroxylase, Esterase, Hydrolase, Laccase, Lipase, No, Oxidized PVA hydrolase, PHA depolymerase, PHB depolymerase, PVA dehydrogenase, Polyurethanase, Polyurethanase A, Polyurethanase B, Polyurethanase esterase, Polyurethane esterase A, Serine hydrolase | HDPE, Impranil, LDPE, Nylon, O-PVA, PBS, PBSA, PCL, PE, PEG, PES, PET, PHA, PHA Blend, PHB, PHBV, PHN, PHO, PLA, PP, PS, PS Blend, PU, PU Blend, PVA, PVC, PVC Blend | 0.04218 |
| OG0000699 | *Pseudomonas* sp. | CP180479 | AP043704, CP020603, CP028517, CP071644, CP071647, CP071649, CP071650, CP071652, CP071655, CP071657, CP071660, CP071661, CP071662, CP071663, CP071664, CP071665, CP071666, CP071667, CP071668, CP071671, CP071674, CP071675, CP071676, CP136718, CP169288, CP178184, **CP180479**, CP180480, CP180481, CP182239, CP187262, CP191382, CP194068, CP194941, CP194945, CP194950, CP195076, CP195082, CP195315, CP195316, CP195762, CP195831, CP195907, CP196684, CP196711, CP196713, CP196739, CP196740, CP196741, CP196749, CP196848, CP196849, CP196919, CP196926, CP196958, CP196969, CP196970, CP197256, CP197353, CP197419, CP198970, CP198974, CP198993, CP198994, CP199001 | Alkane hydroxylase, Esterase, Hydrolase, Laccase, Lipase, No, Oxidized PVA hydrolase, PHA depolymerase, PHB depolymerase, PVA dehydrogenase, Polyurethanase, Polyurethanase A, Polyurethanase B, Polyurethanase esterase, Polyurethane esterase A, Protease, Serine hydrolase | HDPE, Impranil, LDPE, Nylon, O-PVA, PBS, PBSA, PCL, PE, PEG, PES, PET, PHA, PHA Blend, PHB, PHBV, PHN, PHO, PLA, PP, PS, PS Blend, PU, PU Blend, PVA, PVC, PVC Blend | 0.03303 |
| OG0002574 | *Pseudomonas aeruginosa* | CP187262 | CP071660, **CP187262**, CP195762, CP196749, CP196969, CP196970, CP198993, CP198994 | Alkane hydroxylase, Esterase, Hydrolase, Laccase, Lipase, No, Oxidized PVA hydrolase, PHB depolymerase, PVA dehydrogenase, Polyurethanase | HDPE, Impranil, LDPE, Nylon, O-PVA, PBS, PBSA, PCL, PE, PEG, PES, PET, PHA Blend, PHB, PHN, PHO, PLA, PP, PS, PS Blend, PU, PU Blend, PVA, PVC, PVC Blend | 0.02135 |
| OG0000855 | *Pseudomonas* sp. | CP180479 | AP043704, CP071644, CP071647, CP071649, CP071650, CP071652, CP071655, CP071657, CP071658, CP071660, CP071661, CP071662, CP071663, CP071664, CP071665, CP071666, CP071667, CP071668, CP071671, CP071674, CP071675, CP071676, CP136718, CP169288, **CP180479**, CP180480, CP180481, CP182239, CP191382, CP194068, CP194945, CP194950, CP195076, CP195082, CP195315, CP195831, CP196684, CP196711, CP196713, CP196739, CP196740, CP196741, CP196757, CP196758, CP196759, CP196760, CP196761, CP196762, CP196763, CP196764, CP196765, CP196766, CP196767, CP196768, CP196769, CP196848, CP196849, CP196913, CP196915, CP196917, CP196919, CP196920, CP196922, CP196923, CP196924, CP196925, CP196926, CP196927, CP196929, CP196958, CP197196, CP197197, CP197198, CP197199, CP197200, CP197201, CP197203, CP197204, CP197256, CP197353, CP197419, CP198974, CP199001 | Alkane hydroxylase, Esterase, Hydrolase, Lipase, No, Oxidized PVA hydrolase, PHA depolymerase, PHB depolymerase, PVA dehydrogenase, Polyurethanase, Polyurethanase A, Polyurethanase B, Polyurethanase esterase, Polyurethane esterase A, Serine hydrolase | HDPE, Impranil, LDPE, Nylon, O-PVA, PBS, PBSA, PCL, PE, PEG, PES, PET, PHA, PHA Blend, PHB, PHBV, PHN, PHO, PLA, PP, PS, PS Blend, PU, PVA, PVC | 0.01985 |
| OG0000605 | *Pseudomonas aeruginosa* | CP180481 | AP043704, CP020603, CP071644, CP071657, CP071660, CP071662, CP071665, CP071667, CP071675, CP180480, **CP180481**, CP182239, CP187262, CP195315, CP195762, CP196749, CP196848, CP196849, CP196969, CP196970, CP197256, CP198974, CP198993 | Alkane hydroxylase, Esterase, Hydrolase, Laccase, Lipase, No, Oxidized PVA hydrolase, PHA depolymerase, PHB depolymerase, PVA dehydrogenase, Polyurethanase | HDPE, Impranil, LDPE, Nylon, O-PVA, PBS, PBSA, PCL, PE, PEG, PES, PET, PHA, PHA Blend, PHB, PHN, PHO, PLA, PP, PS, PS Blend, PU, PU Blend, PVA, PVC, PVC Blend | 0.01491 |
| OG0002794 | *Pseudomonas aeruginosa* | CP187262 | CP020603, CP187262, CP195082, CP195762, CP195907, CP196749, CP196969, CP198993, CP198994 | Esterase, Laccase, Lipase, No, PHB depolymerase, Polyurethanase A, Polyurethanase B | HDPE, LDPE, PBSA, PE, PES, PET, PHB, PS, PU, PU Blend, PVC Blend | 0.01381 |
| OG0006230 | *Pseudomonas aeruginosa* | CP180479 | CP020603, **CP180479**, CP187262, CP195082, CP195762, CP195907, CP196749, CP196969, CP196970, CP198993, CP198994 | Esterase, Laccase, Lipase, No, PHB depolymerase, Polyurethanase A, Polyurethanase B | HDPE, LDPE, PBSA, PE, PES, PET, PHB, PS, PU, PU Blend, PVC Blend | 0.01287 |

